# The first homosporous lycophyte genome revealed the association between the dynamic accumulation of LTR-RTs and genome size variation

**DOI:** 10.1101/2022.12.06.519249

**Authors:** Ji-Gao Yu, Jun-Yong Tang, Ran Wei, Mei-Fang Lan, Rui-Chen Xiang, Qiao-Ping Xiang, Xian-Chun Zhang

**Affiliations:** State Key Laboratory of Systematic and Evolutionary Botany, Institute of Botany, the Chinese Academy of Sciences, Beijing 100093, China; University of Chinese Academy of Sciences, Beijing 100049, China

**Author notes:** These authors contributed equally to this work (Ji-Gao Yu, Jun-Yong Tang).

**Keywords:** homosporous lycophyte, genome size, LTR-RTs, whole genome duplication, huperzine A

## Abstract

Lycophytes and euphyllophytes (ferns and seed plants) are the two surviving lineages of vascular plants. The modern lycophytes (clubmosses) are herbaceous found either heterosporous (Isoetales and Selaginellales) or homosporous (Lycopodiales). The contrasting genome size between homosporous and heterosporous plants has long been an attractive topic. Most clubmosses are the resource plants of Huperzine A (HupA) which is invaluable for treating Alzheimer’s disease, but the evolutionary trajectory of which in land plants is unexplored. To better understand these fundamental questions, the genome data of a homosporous lycophyte is urgently required. We generated the *Lycopodium clavatum* L. genome by applying a reformed pipeline for filtering out non-plant sequences. The obtained genome size is 2.30 Gb, distinguished in more than 85% repetitive elements of which 62% is LTR. Two whole genome duplications (WGDs) are rigorously detected. The content of LTR-RTs was more than ten times higher in homosporous lycophytes than heterosporous ones, although most appeared within one Mya. Then, we find that the LTR-RTs’ birth-death mode (a much greater birth and extremely slower death) contributes the accumulation of LTR-RTs resulting homosporous lycophyte genome expansion, while in heterosporous lycophytes, the mode is exactly the opposite. Furthermore, the five necessary enzymes of the HupA biosynthetic pathway were identified in the *L. clavatum* genome, but absent in the other land plants. This decoded genome data will be a key cornerstone to elucidating the fundamental aspects of lycophyte biology and land plant evolution.

## INTRODUCTION

Lycophyte represents a sister lineage to the euphyllophytes (ferns and seed plants) (Banks et al., 2011; Bateman, 1996; Gao et al., 2010; Schneider et al., 2009), which is the most ancient and ecologically prominent vascular lineages of the Earth’s terrestrial flora (Schuettpelz et al., 2016; Sessa & Der, 2016), dating back to 392-451 million years ago (Mya) (Barba-Montoya et al., 2018). This systematically distinguished plant group comprises heterospory Isoetales and Selaginellales, and homospory Lycopodiales. The latter lineage is an invaluable medicinal resource for the treatment of Alzheimer’s disease (AD) because the plants contain an important lycopodium alkaloid, Huperzine A (HupA), a potent and reversible acetylcholinesterase inhibitor (Tang et al., 1986; Tang et al., 1989; Yang et al., 2017). Previous studies identified five types of enzymes involved in the biosynthetic pathway of HupA in Lycopodiales, that is lysine decarboxylase (*LDC*) (Bunsupa et al., 2016; Xu et al., 2017), copper amine oxidase (*CAO*) (Sun et al., 2012), type III polyketide synthase (*PKS*) (Wanibuchi et al., 2007), berberine bridge enzyme (*BBE*) and secologanin synthase (*SLS*) (Yang et al., 2017). Homosporous lycophytes are always abundant with many kinds of endophytic fungi (Horn et al., 2013; U’ Ren et al., 2012), some of which produce HupA (Kang et al., 2019). However, the notorious endophytic fungi may hamper the genome sequencing and assembly of homosporous lycophytes.

The striking genome size contrast between homosporous and heterosporous plants has long been an attractive topic. The seed-free vascular plants (lycophytes and ferns), are distinct in having experienced independent homo-heterospory transitions (Szövényi et al., 2021), and the genome size of homosporous lineages tends to be much larger than that of the heterosporous ones in lycophytes and ferns respectively (Kuo & Li, 2019). Previous studies recognized a significant positive relationship between genome size and chromosome number in lycophytes and ferns, but not in the other vascular plants, suggesting that the chromosome architecture of seed-free vascular plants is more static than that in seed-bearing plants (Barker & Wolf 2010; Nakazato et al., 2008; Sessa & Der, 2016; Szövényi et al., 2021). These observations lend support to the longstanding hypothesis that the evolution of genome size in seed-free vascular plants is mainly shaped by the variation of chromosome number mainly resulting from whole genome duplications (WGDs) (Klekowski & Baker, 1966; Haufler, 1987; Huang et al., 2020; Wagner et al., 1979; Wang et al., 2022). However, the genome silencing and rearrangement after recurrent WGDs are contradictory to the relative stasis of chromosomes in seed-free vascular plants, suggesting other processes involved in the evolution of genome size. Another major player contributing to the genome size variation is transposable elements (TEs), especially long terminal repeat retrotransposons (LTR-RTs) (Szövényi et al., 2021). This hypothesis, although widely accepted in seed plants, remains controversial in different studies on seed-free vascular plants (lycophytes and ferns) (Feschotte et al., 2002; Hanson & Leitch, 2002; Wendel et al., 2016; Zedek et al., 2010). Some studies found that there was no correlation between the genome size and the LTR-RTs richness because LTR-RTs activity was too recent to effect (Banks et al., 2011; Wolf et al., 2015; VanBuren et al., 2018). However, Baniaga & Barker (Baniaga & Barker, 2019) found that the timing of median LTR activity was positively correlated with genome size in fern and lycophyte taxa. More recently, a negative correlation between the LTR-RTs insertion time and genome size was reported in angiosperm, and competition between LTR-RTs insertion and deletion in the genome was proposed to interpret the phenomenon (Wang et al., 2021). Based on the recently reported genome data of *Ceratopteris richardii*, and *Adiantum capillus-veneris*, the richness of LTR-RTs was proposed as the main factor affecting genome expansion in homosporous ferns (Fang et al., 2022; Marchant et al., 2019 & 2022). However, neither explorations on the processes underlying the LTR-RTs accumulation nor comparison studies between homosporous and heterosporous seed-free vascular plants have been conducted. The homosporous and heterosporous lycophytes are sister groups with relatively similar genetic backgrounds, which provide a suitable model system to test the hypothesis on LTR-RTs accumulation and further explore the mechanisms underlying the genome size evolution. Four genomes of heterosporous lycophytes (i.e., *Isoetes taiwanensis, Selaginella moellendorffii, S. lepidophylla*, and *S. tamariscina*) have been published (Banks et al., 2011; VanBuren et al., 2018; Wickell et al., 2021; Xu et al., 2018), but no homosporous lycophyte genome is available currently. Considering the crucial phylogenetic position and the medicinal importance of homosporous lycophytes, it is necessary to carry out a genomic study.

In this study, we performed the first homosporous lycophyte genome analyses of *Lycopodium clavatum* L. using high-throughput sequencing technology including both PacBio and Illumina, and proposed a reformed pipeline for decontamination. We subsequently revealed the processes underlying the LTR-RTs accumulation and the association with genome size increase in the homosporous lycophyte genome. We further explored the evolutionary trajectory of the HupA biosynthetic pathway in land plants which laid the foundation for bioprospecting Huperzine and utilization of these valuable plants.

## RESULTS

### Genome assembly

We sequenced the first homosporous lycophyte genome of *L. clavatum* L. using a combination of Illumina and Pacbio high-throughput sequencing strategies. We generated 176.13 Gb (76× in depth) clean reads from Illumina and 235.46 Gb (102× in depth) subreads from Pacbio sequencing platforms (Tables S1 and S2). We initially analyzed the genome by K-mer=17, and the estimated heterozygosity and repetitive sequences in the genome were 0.89% and 72.92%, respectively, indicating an extraordinarily complex genome (Table S3). The genome size of *L. clavatum* was estimated to be 2.53 GB (Table S3, Figure S1), slightly smaller than the DNA content of 2.86 pg/C by flow cytometry (Hanson & Leitch, 2002). We totally assembled 7,228 contigs, and after decontamination (see Method, Figure 1A), we got an accurate *L. clavatum* genome sequence with 7,102 contigs, 2.30 Gb altogether, and contig N50 length is about 1.01 Mb (Figure 2A, Table 1).

**Table 1.** Statistics of the *Lycopodium clavatum* genome.

|  | Type | Number |
| --- | --- | --- |
| Assembly | Total Number of Contigs | 7,102 |
|  | length of Contigs (bp) | 2,304,742,692 |
|  | Shortest Contigs (bp) | 3,171 |
|  | Longest Contigs (bp) | 5,705,076 |
|  | N50 of Contigs (bp) | 1,012,737 (648 Contigs) |
|  | N95 of Contigs (bp) | 85,926 (3,646 Contigs) |
|  | GC content (%) | 38.42 |
| Annotation | Number of protein-coding genes | 23,292 |
|  | % of genes annotated | 87.47% (20,374) |
|  | % of genes with GO terms | 46.66% (10,868) |
|  | Mean gene length (bp) | 13,019.30 |
|  | Mean exon length (bp) | 420.33 |
|  | Exon length (bp) | 50,464,920 |
|  | Intron length (bp) | 252,780,510 |
|  | Number of pseudogenes | 5,771 |
|  | Length of pseudogenes (bp) | 11,692,858 |
|  | Mean pseudogene length (bp) | 2026.14 |

**Figure 1.**
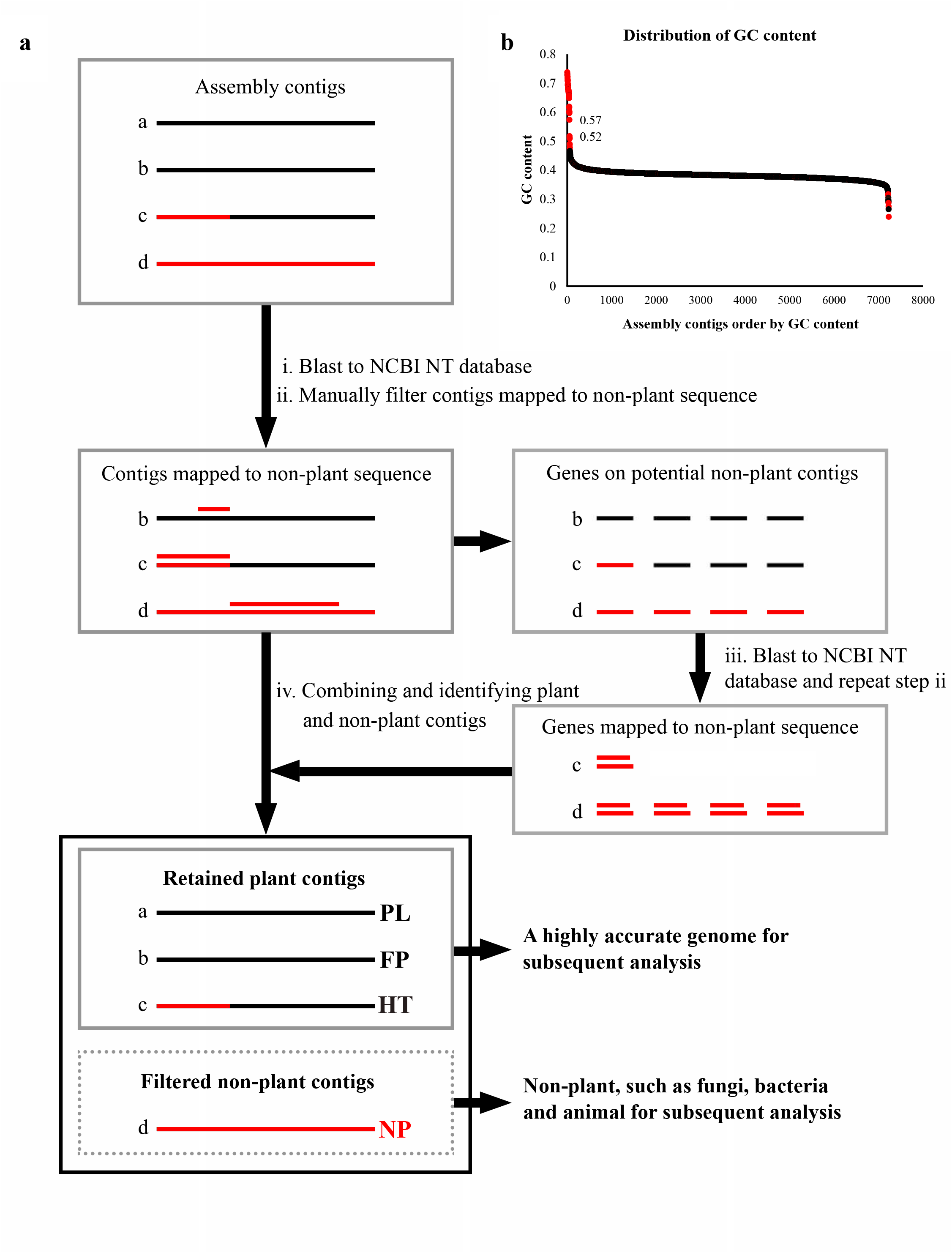
Workflow of the post-assembly decontamination pipeline. **a**, It includes six steps: (i) Blast to NCBI NT database; (ii) Manually filter the non-plant alignment; (iii) Blast to NCBI NT database and repeat step ii; (iv) Combining and identifying the contigs of plant and non-plant. PL, plant; FP, false positive; HT, horizontal transfer; NP, non-plant. The red line represents the non-plant sequence, and the black line represents the plant sequence. **b**, The red dots represent the GC content of non-plant contigs determined by this pipeline. The detailed information for GC content data in this graph was listed in Table S15.

**Figure 2.**
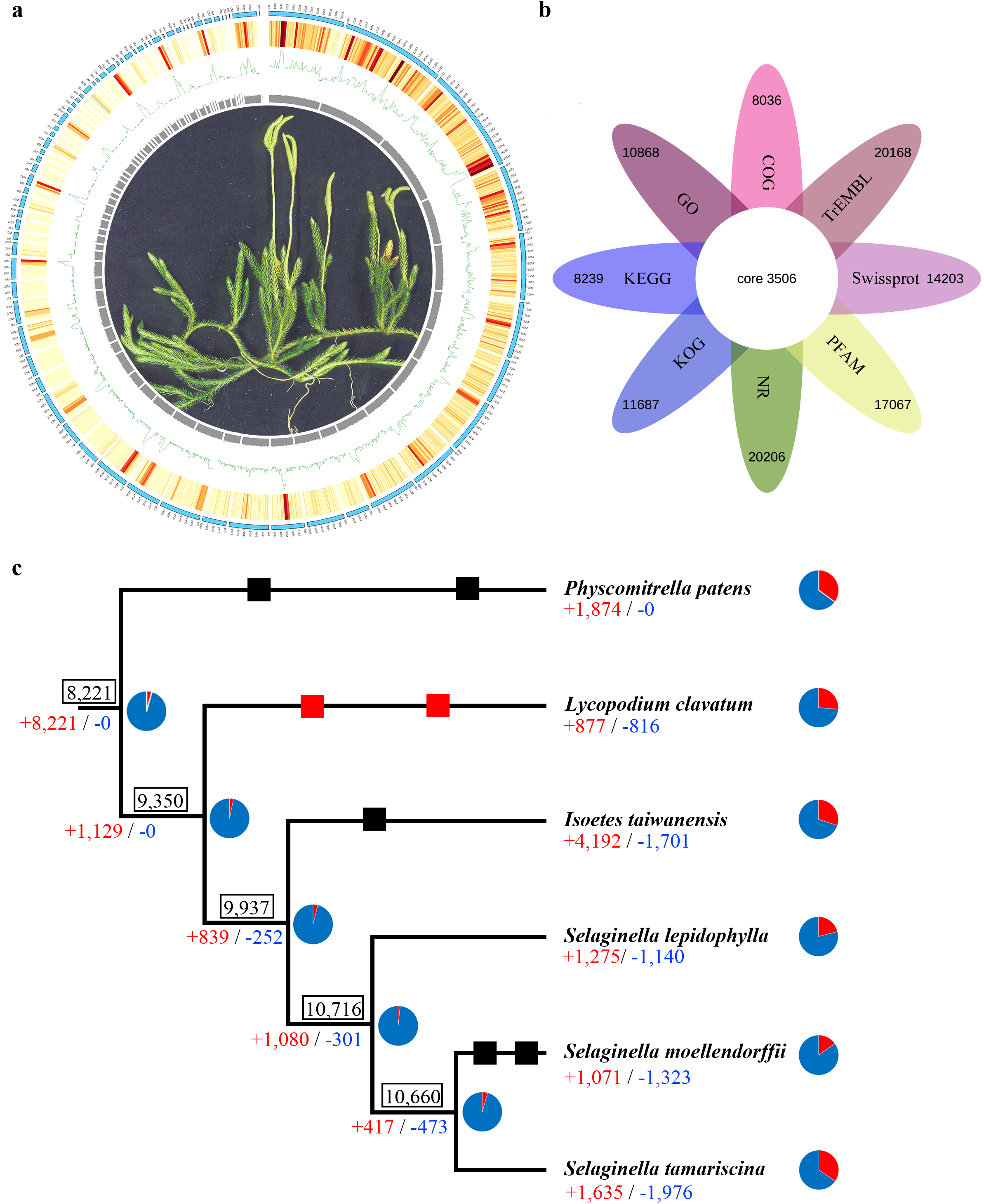
Genome features and evolutionary analysis of the *Lycopodium clavatum* genome. **a**, Genome features depicted by 72 groups arranged by 7,102 contigs in ASCII order and divided by 100 contigs. From outer to inner, the karyotype bands, gene density, long terminal repeat (LTR) density, and GC content density. Center: the phenotype of *L. clavatum*. **b**, Venn diagram illustrating the number of annotated genes by eight databases, including non-redundant protein sequence database (NR), cluster of orthologous groups of proteins (COG), gene ontology (GO), Kyoto encyclopedia of genes and genomes (KEGG), clusters of orthologous groups for eukaryotic complete genomes (KOG), translated EMBL nucleotide sequence data library (TrEMBL), PFAM and Swissport databases. **c**, Gain (+) and loss (-) of gene families among six plant species. The numbers of gained (red) and lost (blue) gene families are shown below the branches. The boxed number indicates the gene-family size at each node. The newly identified WGDs (red rectangles) and published WGDs (black rectangles) are shown on the branches. The pie charts show the proportion of duplicated genes in each node and plant species.

Different strategies were used to assess the quality and completeness of the *L. clavatum* genome. First, 84.60% of Illumina paired-end reads were successfully mapped to the assembly genome, and all mapped reads accounted for 93.79% (Table S4). Additionally, core eukaryotic gene mapping approach (CEGMA) analysis indicated that 73.39% (338) of core eukaryotic genes were detected in the *L. clavatum* genome (Table S5). Then, Benchmarking Universal Single-Copy (BUSCO) analysis showed that 75.46% of the 1,614 core embryophyte genes were present in the *L. clavatum* genome, and 69.33% of them were single-copy genes (Table S6). We also used the LTR Assembly Index (LAI) to evaluate the assembly quality (Ou et al., 2018), and the LAI score of *L. clavatum* genome is 20.27 (Table S7).

### Genome annotation

We used multiple *de novo* and homology-based prediction procedures to annotate repetitive elements. An extraordinarily high proportion (85.32%, 1.97 Gb) in the *L. clavatum* genome was identified as repetitive elements, including Class I (retrotransposons, 76.42%), Class II (DNA transposons, 7.71%), potential host genes (0.37%), simple sequence repeats (0.29%), and unclassified elements (5.32%) (Table S8). Copia and Gypsy superfamily of LTR accounted for 36.25% and 26.09%, respectively.

The gene model prediction combined with homologous strategy, ab initio strategy, and transcript evidence-supported strategy. A total of 23,292 genes (distributed in 3,775 contigs) were predicted, the average gene and exon lengths were 13,019.30 bp and 420.33 bp, respectively (Figure 2b, Table 1 and S9). (Non-redundant protein sequences, NR) homologous species distribution exhibited that the *L. clavatum* genome was the closest to that of *S. moellendorffii* (6,684, 32.81%), followed by that of *Physcomitrella patens* (4,386, 21.53%) (Table S10, Figure S2). Gene function enrichments of GO mostly focused on ATP binding, membrane, and nucleus (Table S11). Gene function enrichments of COG and KOG mostly focused on replication, recombination, repair, and posttranslational modification, protein turnover, and chaperones (Tables S12 and S13, Figures S3 and S4,). Furthermore, 4,567 tRNAs, 530 rRNAs, 24 miRNAs, and 5,771 pseudogenes were predicted in the *L. clavatum* genome (Table S14).

### Discovery and removal of non-plant sequences

The total GC content of the *L. clavatum* genome is 38.42% (Table 1), while some contigs exhibit abnormally high GC content. Interestingly, an obvious breakpoint was found between 0.52 and 0.57 (Figure 1b, Table S15). Forty-nine contigs with GC content higher than 0.52 were compared with the Nucleotide Sequence Database (NT) in National Center for Biotechnology Information (NCBI), and we found that these contigs only hit non-plant species, such as fungi, bacteria, and other distant species (-max_target_seqs 1, -evalue 1e-5) (Table S16). Furthermore, in order to verify the reliability of high GC content as non-plant sequences, we calculated the GC content of alignment regions (non-plant sequences) and non-alignment regions (plant sequences) in the contigs with abnormal GC content, and there was no difference for the GC content of the alignment region and the non-alignment region in the same contig (Table S17). Therefore, non-plant sequences and *L. clavatum* genome sequences could not be separated just by the abnormal GC content.

We assumed that non-plant sequences and the *L. clavatum* genome sequences were unable to be assembled into one contig, and horizontal gene transfer could be assembled into the *L. clavatum* genome sequences. We further compared all 7,228 contigs with the NCBI NT database and found that 131 contigs (including 49 contigs with abnormal GC content) were the best alignment for non-plant species which were regarded as the potential non-plant contigs. Then, we compared 601 genes on the 131 contigs with the NT database and found one contig with horizontal transfer genes and five false positive contigs which only mapped to plant gene sequences. Eventually, we got 126 contigs as non-plant sequences with a total length of 6.56 Mb (0.28% of the whole genome) and 585 genes (in 99 contigs) (Table S18). Function enrichment of the non-plant genes mainly involving in energy production and transmembrane transport mediated by ATP-binding cassette transporters (Tables S19 and S20), which indicates that the endophytes may provide the carbon sources to the gametophytes of Lycopodiales (Horn et al., 2013; Szövényi et al., 2021).

### Comparative analyses of gene family

We investigated the gene families in the early diverged land plants including *P. patens, L. clavatum, I. taiwanensis, S. lepidophylla, S. moellendorffii*, and *S. tamariscina*. We clustered 22,610 orthogroups, accounting for 84% of all six genomes (Table S21), 8,534 orthogroups of *L. clavatum* shared with other plants, and 877 species-specific orthogroups appeared to be unique in *L. clavatum* (Table S22). The unique genes in *L. clavatum* were enriched in posttranslational modification, protein turnover, chaperones, secondary metabolites biosynthesis, transport and catabolism, and transcription (Tables S23 and S24).

Comparative genomics exhibited that the common ancestor of lycophytes had 9,350 gene families. For the *L. clavatum* genome, it showed the least loss and the least gain from the common ancestor (Figure 2c). The heterosporous lycophyte ancestor included 9,937 gene families, with only a small number of gain (839) and minimal loss (252) during the evolution from the common lycophyte ancestor node to the heterosporous lycophyte node (Figure 2c). **The processes of LTR-RTs’ accumulation** It has long been recognized that the abundance of LTR-RTs is positively correlated with the genome size, which is also applicable to lycophytes. There are three types of LTR-RTs: intact LTR-RTs (I, with complete *Gag*-*Pol* protein sequence), solo-LTR-RTs (S, LTR-RTs paralogs that lack any *Gag*-*Pol* homologs in both the upstream and downstream sequences) and truncated LTR-RTs (T, LTR-RTs with *Gag*-*Pol* sequences on one side of flanking sequences), and *S:I* values indicate the deletion rate of LTR-RTs which are particularly prone to removal when the value is higher than 3 (Lyu et al., 2018). We identified all of three types LTR-RTs in the *L. clavatum* genome and three published heterosporous lycophyte genomes (*I. taiwanensis, S. tamariscina*, and *S. moellendorffii*). For homosporous lycophyte, it exhibited a much greater birth (I+S+T) (Lyu et al., 2018) and extremely slower death (*S:I*<1) model of LTR-RTs, resulting in the largest accumulation of LTR-RTs and expansion of the *L. clavatum* genome (Figure 3ab, Table S25). For heterosporous lycophytes, all have a lower birth and faster death (*S:I*, 2.81 to 5.60) model of LTR-RTs, resulting in a small accumulation of LTR-RTs and relatively small genomes in heterosporous lycophytes (Figure 3ab, Table S25). We detected a positive correlation between LTR-RTs and the genome sizes in lycophytes (Figure S5). All the three types of LTR-RTs exhibited a positive correlation coefficient with the genome sizes (R^2^ = 0.98, 0.68,0.67; F-test, P-value = 1e-3, 3.08e-5, 9.03e-4), while *S:I* ratios showed a negative correlation with the genome sizes (R^2^ = 0.29; F-test, P-value = 1.43e-8, Figure S5).

**Figure 3.**
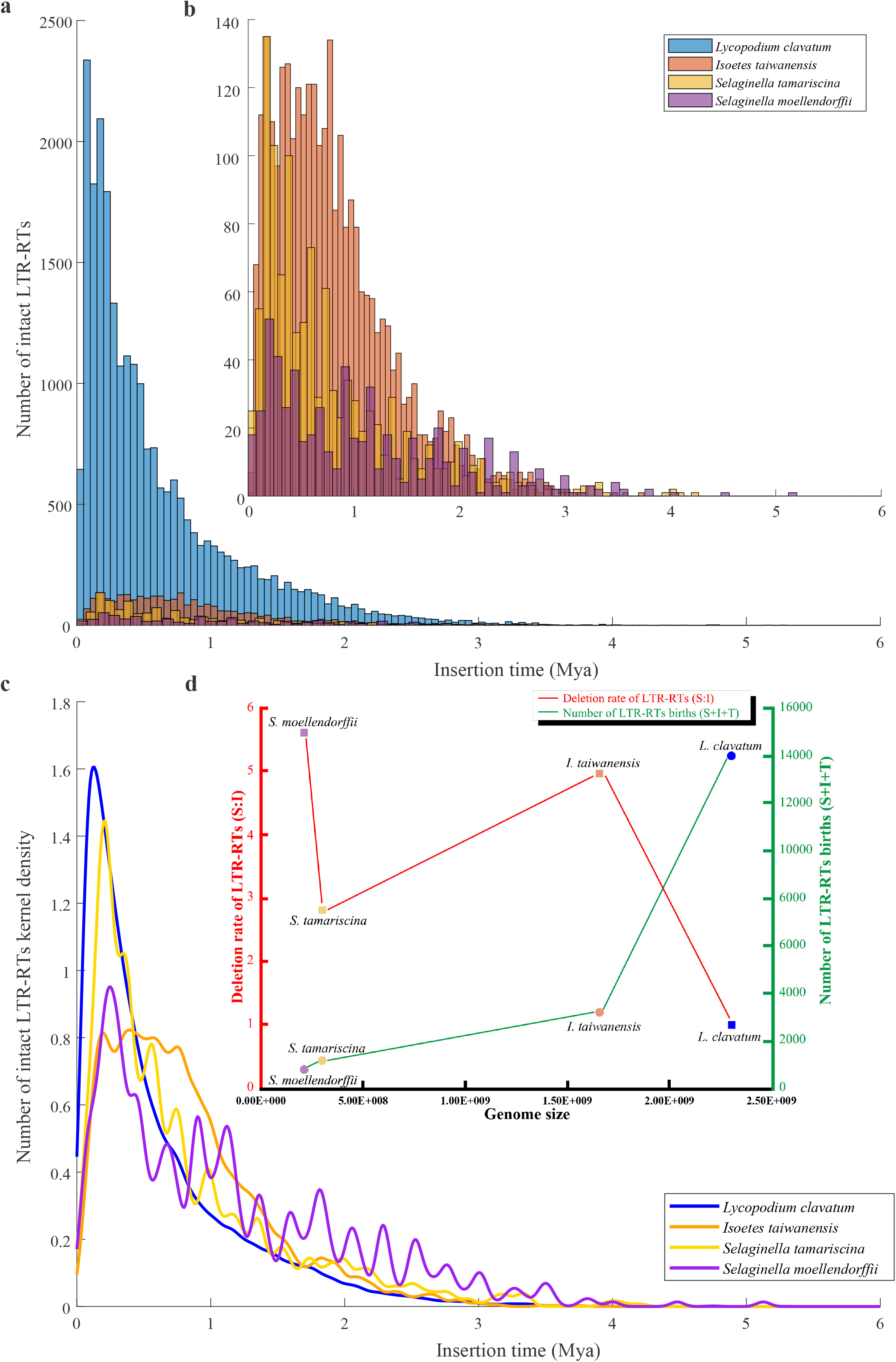
Distribution of intact LTR-RT insertion time in lycophytes. **a**, Distribution of intact LTR-RT insertion time in lycophytes. **b**, Distribution of intact LTR-RT insertion time in lycophytes except *L. clavatum*. **c**, Fitting curve of distribution of LTR-RT insertion time in lycophytes. **d**, Genome size, deletion rate and birth of LTR-RTs in lycophytes.

In addition, we examined the insertion time of intact LTR-RTs. We found that the insertion time of most LTR-RTs in lycophytes is very young (< 1 Mya) (Figure 3c). For homosporous lycophyte *L. clavatum*, the concentrated outbreak of LTR-RTs was around 0.11 Mya, and the insertion of 80.68% LTR-RTs was in one Mya; For heterosporous lycophyte, *I. taiwanensis*, it had a continuous LTR-RTs insertion in 0.16-1 Mya, and the number of LTR-RTs during this period accounted for 70.76%; For another heterosporous *Selaginella*, LTR-RTs showed the trend of multiple insertions in different periods during six Mya. The highest peak of insertion time was 0.20 Mya in *S. tamariscina*, which was 0.25 Mya later than that of in *S. moellendorffii* (Figure 3c). Furthermore, we found a sharp contrast of LTR-RTs birth and death model (Figure 3d). For homosporous lycophyte, it experienced a much greater birth and extremely slower death of LTR-RTs, while the heterosporous lycophytes possessed a lower birth and faster death model (Figure 3d).

### Two specific whole-genome duplications were detected in the *Lycopodium clavatum* genome

Using a joint strategy of synonymous substitutions per synonymous site (Ks) and 8,498 phylogenetic trees, we identified at least two ancient whole-genome duplications (WGDs) in the *L. clavatum* genome. First, we tried the syntenic analysis which is the most direct and powerful gold standard for identifying WGDs (Jiao et al., 2914; Wang et al., 2018), but it is difficult to find the collinearity of the *L. clavatum* genome (Figure S6). Actually, we only identified 100 WGD gene pairs (197 genes) in 18 syntenic genomic segments and 16,256 dispersed duplicated gene pairs (18,778 genes) using DupGen_finder (Tables S26 and S27) (Qiao et al., 2019). A large number of dispersed duplicated genes and a small number of WGD genes suggest that there may be polyploidy in *L. clavatum* genome. Based on the analysis of 8,498 low-copy homologous gene trees by tree2gd (https://sourceforge.net/projects/tree2gd/), we found that the proportion of duplicated genes reached 35.69%, which indicated that there were WGDs in *L. clavatum* (Figure 2c, Table S28).

In addition, we made Ks statistics for the two best-matched gene pairs produced by Blastp, removed the gene pairs on the same contig (eliminate the influence of tandem repeat genes as far as possible), and finally obtained 19,583 gene pairs, a total of 17,971 genes, distributed on 3,570 contigs. Two obvious peaks were detected (Table S29, Figure S7), and Ks values of the two peaks (alpha and beta) were 0.45 (σ = 0.14) and 1.27 (σ = 0.67), respectively (Figure 4a, Table S30), revealing that there were two WGDs in *L. clavatum* genome (Figure 2c).

**Figure 4.**
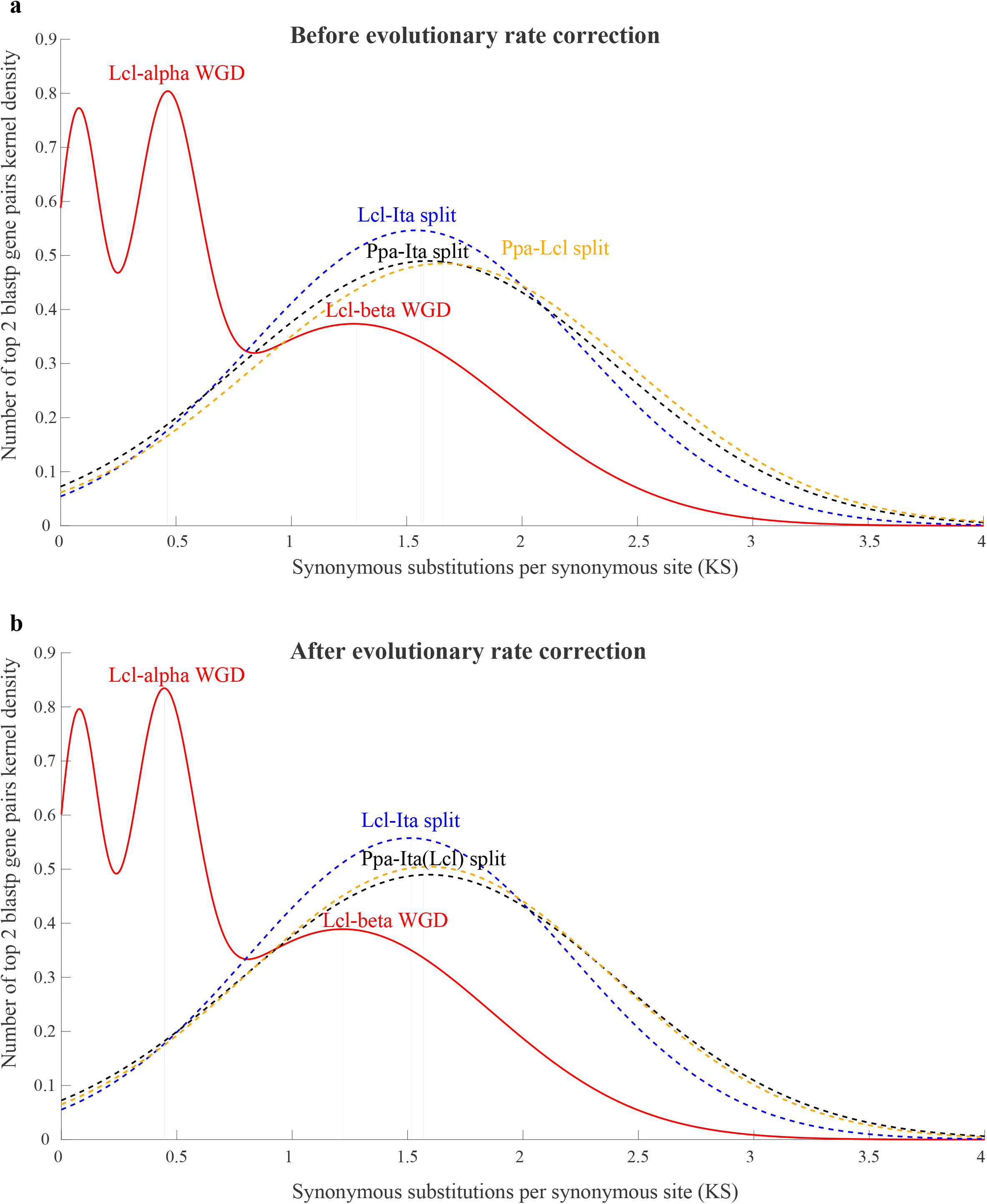
Distribution of synonymous nucleotide substitutions (Ks) for top two best-matched pairs of Blastp in *Physcomitrella patens* (Ppa), *Lycopodium clavatum* (Lcl), and *Isoetes taiwanensis* (Ita). **a**, Distribution of Ks (top two best-matched Blastp pairs) within the *L. clavatum* genome (solid curves) and between two species (dashed curves) before evolutionary rate correction. **b**, Fitting curve of distribution of Ks (top two best matched Blastp pairs) within *L. clavatum* genome (solid curves) and between two species (dashed curves) after evolutionary rate correction.

In order to infer the relative position of the two WGDs, we corrected the evolutionary rate with *P. patens* as the outgroup. The Ks peaks of differentiation between *P. patens* and *L. clavatum, P. patens* and *I. taiwanensis*, and *L. clavatum* and *I. taiwanensis* were 1.66 (σ = 0.82), 1.59 (σ = 0.81) and 1.54 (σ = 0.72) respectively (Figure 4a, Table S30). The evolution rate of *L. clavatum* was 4.15% which was faster than that of *I. taiwanensis*. After correction, the alpha and beta Ks peaks of *L. clavatum* were 0.44 (σ=0.14) and 1.22 (σ=0.65), respectively, and the Ks peak of differentiation between *L. clavatum* and *I. taiwanensis* was 1.51 (σ= 0.70) (Figure 4b, Table S30). Therefore, the two specific WGDs occurred after the differentiation between *L. clavatum* and *I. taiwanensis* and affected the formation of *L. clavatum* genome.

### Pathway evolution of Huperzine A biosynthesis in land plants

It is known that many species of Lycopodiaceae yield HupA. To detect the integrality of the function module of HupA biosynthesis in *L. clavatum* (Figure 5a), we analyzed the key enzyme genes known to involve in HupA biosynthesis in the genomes of *L. clavatum* and other six species (*P. patens, I. taiwanensis, S. moellendorffii, S. tamariscina, S. lepidophylla*, and *Arabidopsis thaliana*). Our result showed that only *L. clavatum* had all five types of enzymes (Figure 5b, Table S31). Although all of the investigated species had the *PKS, BBE*, and *SLS*, both *P. patens* and *A. thaliana* were short of *LDC* which restricts the first step of the biosynthetic route, suggesting the acquisition of *LDC* occurred in the ancestor of lycophytes; *I. taiwanensis* and other three *Selaginella* species lacked *CAO* which restricts the second biosynthetic step, indicating *CAO* has been lost in the ancestor of heterosporous lycophytes. These results indicated that the HupA biosynthetic pathway in homosporous lycophyte was most complete compared with moss and the other vascular plants including the heterosporous lycophytes. The key innovation of the pathway evolution in homosporous lycophytes might be crucial for HupA biosynthesis.

**Figure 5.**
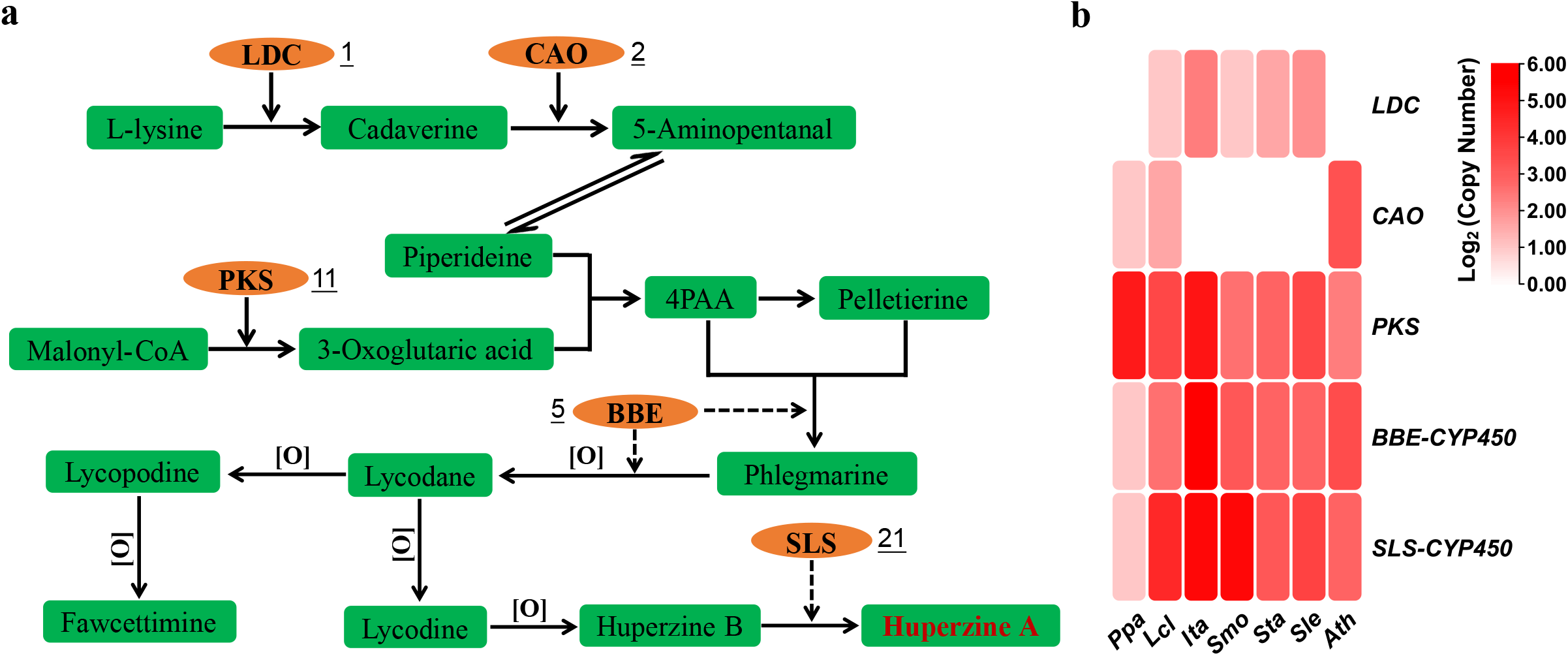
The biosynthetic pathway of Huperzine A and the key enzymes in *Lycopodium clavatum*. **a**, The proposed biosynthetic pathway of Huperzine A modified from a previous study (Yang et al., 2017). The green boxes represent the mesostate that involves in the biosynthetic process. The orange ovals represent the key enzymes, the underline numbers indicate the copy number of the enzyme genes. LDC, CAO, PKS, BBE, and SLS respectively refer to lysine decarboxylase, copper amine oxidase, type III polyketide synthase, berberine bridge enzyme, and secologanin synthase, 4PAA refers to 4-(2-piperidyl) acetoacetate. **b**, The heatmap shows the copy number pattern of the enzyme genes in different species. *Ppa, Lcl, Ita, Smo, Sta, Sle*, and *Ath* refer to *Physcomitrella patens, Lycopodium clavatum, Isoetes taiwanensis, Selaginella moellendorffii, Selaginella tamariscina, Selaginella lepidophylla*, and *Arabidopsis thaliana*, respectively.

## DISCUSSION

### Genome characters of the first homosporous lycophyte, *Lycopodium clavatum* L

The genome of *L. clavatum* obtained here, as the first homosporous lycophyte data resource, fills the last and foremost information gap in vascular plants. This homosporous lycophyte not only has extremely high repetitive sequences (more than 85%) in the genome (Table 1), but also the average gene length (13,019 bp) is much longer than that of the heterosporous lycophytes (the longest is 2,917 bp in *I. taiwanensis*, the shortest is 1,915 bp in *S. tamariscina*) (Banks et al., 2011; VanBuren et al., 2018; Wickell et al., 2021; Xu et al., 2018). The homosporous ferns also have similar features, that is, *C. richardii* and *A. capillus-veneris*, both of which have more than 85% repetitive sequences, and the average length of genes has reached 14,457 bp and 23,212 bp respectively (Fang et al., 2022; Marchant et al., 2022). Different from the homosporous ferns (heterozygosity is 0.25%) (Fang et al., 2022), the homosporous lycophyte has high heterozygosity reaching 0.89% (Table S3). The combination of high repetition and high heterozygosity reflects the extraordinary genome complexity and results in a fragmented genome assembly (Table 1), which hampers further exploration of the genome structure and the evolution in lycophytes and vascular plants. Using Ks-based and phylogenomic-based methods, we both identified two independent WGD events in the *L. clavatum* genome, which is consistent with previous studies using transcriptome data (One Thousand Plant Transcriptomes Initiative, 2019; Xia et al., 2022). Furthermore, the least loss and the least gain of gene families in the homosporous lycophyte genome (Figure 2c), are of great significance for studying the evolution of the common ancestor of lycophytes and euphyllophytes.

Many species of homosporous lycophytes are medicinal plants. Some of them are famous for producing HupA which is effective in the treatment of AD (Chen et al., 2021; Tang et al., 1986; Yang et al., 2017). Five important enzymes (*LDC, CAO, PKS, BBE*, and *SLS*) have been proven to be involved in the biosynthetic pathway of HupA in several studies using transcriptomic data in the homosporous lycophyte genera *Huperzia* and *Phlegmariurus* (Luo et al., 2010a, b; Yang et al., 2017). By comparison analysis of the genome data of homosporous lycophytes, moss, and representatives of other vascular plants including the heterosporous lycophytes, we found that the HupA biosynthetic pathway in homosporous lycophytes was the most complete (Figure 5b). Although the *L. clavatum* genome contains all five key enzymes for HupA synthesis, no HupA was reported in *L. clavatum* plants previously (Ainge et al., 2002; Sahidan et al., 2012). It indicates that some other key enzymes are not discovered yet, and further study is needed on the biosynthetic pathway of HupA. Anyway, it is a key genomic innovation of homosporous lycophyte, which lays the foundation for bioprospecting HupA and sustainable utilization of these valuable plants (Kang et al., 2019; Yang et al., 2017).

### The dynamic of LTR-RTs accumulation underlying the genome size increase in the homosporous lycophyte

Lycophytes, similar to ferns, have experienced homo-heterospory transitions (Szövényi et al., 2021). Most studies showed that homosporous lineages have a much larger genome size than the heterosporous group (Fang et al., 2022; Huang et al., 2020; Kuo & Li, 2019; Lang et al., 2018; Marchant et al., 2022; Wickell et al., 2021), and the abundance of LTR-RTs is regarded as a major factor in genome expansion (Fang et al., 2022; Marchant et al., 2019 & 2022).

However, the processes underlying the LTR-RTs accumulation remain unexplored yet. Compared with heterosporous lycophytes, the LTR accumulation in homosporous lycophyte was dozens of times higher, although most LTR-RTs were generated within one Mya (Figure 3). Furthermore, we found a sharp contrasting model of LTR-RTs’ birth and death between homosporous and heterosporous lycophytes contributes to the accumulation and removal of LTR-RTs resulting in genome size variation (Figure 3d), rather than the timing of median LTR-RTs (Baniaga & Barker, 2019). For homosporous lycophyte, it experiences a much greater birth and extremely slower death of LTR-RTs, while the heterosporous lycophytes possesses a lower birth and faster death model (Figure3d). Combined with the super high proportion of LTR (62%) in homosporous lycophyte *L. clavatum* genome, we propose that the accumulation of LTR-RTs has an important contribution to the genome expansion of homosporous lycophyte. Considering the proportion of LTR ranges from 15.72% (in *I. taiwanensis*) to 35.6% (in *S. moellendorffii*) (Banks et al., 2011; Wickell et al., 2021; Xu et al., 2018), this pattern indicates that the removal of LTR-RTs may have contributed to the relatively small genomes of heterosporous lycophytes.

In addition, TEs insertion of the intron is believed to be the major contributor to the expansion of gene length in *C. richardii* (Marchant et al., 2022), but the total gene length, including introns, only accounts for 7.14% of the whole genome, which is too small to have an effect on the expansion of the genome. Similarly, the homosporous lycophyte genome also has long-length genes (average 13,019 bp), but the total gene length accounts for 13.16% of the whole genome, which is little impact on the genome expansion as well.

In brief, our conclusions support the contribution of LTR richness to genome size evolution, and further explore the mechanism of LTR accumulation, that is the difference between LTR in birth and death. But the causes of the difference in LTR birth and death patterns between homosporous and heterosporous lycophytes remain poorly understood. It may be due to the difference in reproductive strategies (Baniaga & Barker, 2019), or other reasons that need to be explored in the future.

### A reformed pipeline for post-assembly decontamination

Endophytic fungi and bacteria are omnipresent and present in most of land plants (including bryophytes and vascular plants), especially in bryophytes (U’ Ren et al., 2012). During genome sequencing, the genomic sequences from these endophytic fungi, bacteria or other distant species are also sequenced which may lead to confounding extraction of sequenced reads for the accuracy of genome assembly, and thus the decontamination process is necessary (Lang et al., 2018; Li & Barker, 2020; Zhang et al., 2020). For the homosporous lycophyte in our study, the genome survey failed to detect the potential impact of contamination on genome assembly (Table S32). However, there is an obvious gap (between 0.52 and 0.57) in the GC content distribution of contigs, indicating that the contamination cannot be ruled out (Figure 1b, Table S15), and such contamination seems to be false sequences of endophytes because of the frequent appearance of endophytic fungi that produce HupA in homosporous lycophytes (Kang et al., 2019; Yang et al., 2017). As expected, all of 49 contigs with abnormally high GC content were mapped to non-plant sequences, such as fungi, bacteria and other distant species, by comparison with NCBI NT database. These contigs should be removed from the plant genome, and the genes on these contigs are easy to be erroneously identified as horizontal gene transfer. However, high GC content cannot be used as an indicator to identify endophytic or contamination sequences, given that no difference were found in GC content between non-plant comparison regions and other regions (regarded as plant regions) on the same contig (Table S17). In order to decontaminate more strictly, based on sequence similarity, we developed a reformed pipeline for post-assembly decontamination and successfully identified one horizontal gene transfer and non-plant sequences, including not only 49 contigs with abnormally high GC content (0.57-0.74), but also other contigs with a GC content ranging from 0.24 to 0.52 (Figure 1b, Table S15). In addition, most HupA-producing endophytic fungi were identified in non-plant sequences using our pipeline (Table S33) (Shu et al., 2014; Su & Yang, 2015; Wang et al., 2011; Zhang et al., 2015). Compared with previous decontamination processes (Lang et al., 2018; Li & Barker, 2020; Zhang et al., 2020), our pipeline also considered the errors caused by horizontal gene transfer and false positives (detail in Method), by combining the evidences of assembly sequence and gene. But there is also a weakness, that is, a lot of manual screening is required, and non-plant components need to be filtered out from the assembled and annotated genomes.

Our post-assembly decontamination pipeline provides an alarm for future plant genome sequencing. Decontamination may be necessary for the whole land plants, even if no pollution is found in the genome survey. Separating the non-plant sequences, such as fungi, bacteria, and other distant sequences from the plant genome to obtain an accurate plant genome, in general considering the symbiotic relationship, is of great significance for plant evolution and the interaction between endophytes and plants.

## EXPERIMENTAL PROCEDURES

### Sequencing and assembly

The materials of *L. clavatum* were collected from the Changbai Mountain Scenic Area, Jilin Province, China. The total genomic DNA of *L. clavatum* was isolated from fresh plant tissues with the CTAB method as described in Li et al., 2013. Three library construction were performed with the NEBNext DNA Library Prep Kit (New England Biolabs, Ipswich, Massachusetts, USA). The sequencing was carried out by Illumina HiSeq 2000 platform (Illumina, San Diego, California, USA) with 150-bp insert libraries. One paired-end (PE) library with an insert size of 150bp was performed with the Dynabeads mRNA DIRECT Kit (invitrogen), and sequenced on the Illumina Nova seq6000 platform. Then, we used Genome Characteristics Estimation (GCE, v1.0.2) to perform genome survey analysis of *L. clavatum* (Liu et al., 2013). For PacBio library construction, genomic DNA was sheared to 40 kb and sequenced on the PacBio Sequel I system. In order to verify whether the data is polluted, we randomly extracted 500,000 reads from Illumina clean reads and compared with the NT database (Table S32). As expected, the sequence data were effective for subsequent genome assembly.

All PacBio clean reads were self-corrected by Canu (Koren et al., 2017), and then assembled with smartdenovo (https://github.com/ruanjue/smartdenovo) based on the corrected data. Then three rounds of correction were performed with Pilon (Walker et al., 2014) using the Illumina data, and then the software purge_haplotigs was used to filter redundancy (Roach et al., 2018).

### Genome quality assessment

The CEGMA (with default parameters) (Parra et al., 2007), BUSCO (with parameter: -l embryophyta_odb10) (Simao et al., 2015), and LTR Assembly Index (LAI, with parameters: -window 500000 -step 50000 -totLTR 61) (Ou et al., 2018) pipelines were used to assess the completeness and accuracy of the genome assembly. In addition, we aligned Illumina paired-end reads to the *L. clavatum* genome and assessed the assembled proportion using BWA and SAMtools (Li & Richard, 2009; Li et al., 2002).

### Repeat annotation

The identification of the transposon element (TE) was based on a combination of homology and de novo predictions. We first customized a de novo repeat library of the genome by RepeatModeler with default parameters (combined with RECON v1.08), RepeatScout, and LtrHarvest) (Bao & Eddy, 2002; Ellinghaus et al., 2008; Flynn et al., 2020; Price et al., 2005). Then full-length long terminal repeat (LTR) and non-redundant LTR library were identified by LTR_retriever with default parameters (Combined with both LTR harvest and LTR_finder) (Ou & Jiang, 2018; Xu et al., 2007). A non-redundant species-specific TE library was constructed by combining the de novo TE sequences library above with the known Repbase (version 19.06), REXdb (V3.0), and Dfam (v3.2) database. Final TE sequences in the *L. clavatum* genome were identified and classified by homology search against the library using RepeatMasker v4.1.2-p1 (Tarailo-Graovac & Chen, 2009). Tandem repeats were annotated by Tandem Repeats Finder (Benson, 1999).

### Gene prediction and annotation

We integrated de novo prediction, homology search, and transcript-based assembly to annotate protein-coding genes in *L. clavatum* genome. The de novo gene models were predicted by the two ab initio gene-prediction software tools, Augustus v2.4 and SNAP v2006-07-28 (Korf et al., 2004; Stanke et al., 2006). For the homolog-based approach, GeMoMa v1.7 was performed by using a reference gene model from one model species *A. thaliana*, and three close species *P. patens, S. moellendorffii*, and *Azolla filiculoides* (Keilwagen et al., 2019). For the transcript-based prediction, RNA-sequencing data were mapped to the reference genome by Hisat v2.0.4 (Kim et al., 2015) and assembled by Stringtie v1.2.3 (Pertea et al., 2015). GeneMarkS-T v5.1 was used for gene prediction based on the assembled transcripts (Tang et al., 2015). The PASA v2.0.2 (Haas et al., 2003) software was used to predict genes based on the unigenes assembled by Trinity v2.11 (Grabherr et al., 2013). Gene models from these different approaches were combined by EVM v1.1.1 (Haas et al., 2008) and updated by PASA. The final gene models were annotated by searching the GenBank Non-Redundant (NR, 20200921), TrEMBL (202005), PFAM (33.1), SwissProt (202005), Cluster of Orthologous Groups of proteins (COG, 20110125), eukaryotic orthologous groups (KOG, 20110125), gene ontology (GO, 20200615) and Kyoto Encyclopedia of Genes and Genomes (KEGG, 20191220) databases.

### Pseudogene prediction

Pseudogene sequences are usually similar to corresponding functional genes but may have lost their biological functions because of genetic mutations, such as insertion and deletion. The GenBlastA v1.0.4 (She et al., 2009) program was used to scan the whole *L. clavatum* genome after masking predicted genes. Putative candidates were analyzed by searching for non-mature mutations and frame-shift mutations using GeneWise v2.4.1 (Birney et al., 2004).

### Noncoding RNAs annotation

Non-coding RNAs are usually divided into several groups, including miRNA, rRNA, tRNA, snoRNA, and snRNA. The tRNAscan-SE v1.3.1(Lowe & Eddy, 1997) was used to predict tRNA with eukaryote parameters, and barrnap v0.9 (Loman, 2017) was used to identify the rRNA genes. The miRNA was identified by searching the miRBase (release 21) database (Griffiths-Jones et al., 2006). The snoRNA and snRNA genes were predicted using INFERNAL against the Rfam (release 12.0) database (Nawrocki & Eddy, 2013).

### Post-assembly decontamination

Considering that non-plant sequences (such as fungi and bacteria) can’t be assembled into the plant genome, but the sequence of horizontal gene transfer and plant genome sequences can be assembled into the same contig, we proposed a reformed pipeline for filtering out non-plant sequences from the plant genome. First, to compare the assembly contigs with the NT database by the blast (-max_target_seqs 1, -evalue 1e-5); second, to filter the best matching assembly contigs with non-plant sequences manually as potential non-plant contigs; third, to compare the annotated genes with the potential non-plant contigs from the second step with NT database, and to repeat the second step afterward; Fourth, if the sequences and genes on the same contig both mapped to the non-plant sequences, then this contig is considered as the final non-plant contig. In the third step, we also considered the affection of false positives and horizontal gene transfer. For the same contig, if the genes can be mapped to non-plant sequences and the other region can be mapped to plant sequences, then the genes are considered as horizontal gene transfer.

### Polyploidy identification

For WGD identification, we used three strategies, synteny-based, Ks-based, and phylogenomic-based methods. DupGen_finder was used to find the collinear region, then the best two matched gene pairs were extracted for calculating the Ks value, and the Nei–Gojobori approach (Nei & Gojobori, 1986) was implemented by the Bioperl Statistical module for Ks-based method. Besides, in order to reduce the influence of tandem repeat genes on polyploidy identification as much as possible, the gene pairs located at the same contig were removed here. The R package gg-plot was used to draw the histogram and Matlab was used to estimate the density of Ks distribution. Then we performed the Gaussian multipeak fitting of the curve by cftool to identify WGDs (Wang et al., 2018). In addition, tree2gd (v1.0.40, https://github.com/Dee-chen/Tree2gd) was used to infer WGDs for the phylogenomic-based method.

### Identification of intact, solo- and truncated LTR-RTs

We customized a de novo repeat library of the genome by RepeatModeler with default parameters (combined with RECON v1.08, RepeatScout, and LtrHarvest) (Ellinghaus et al., 2008; Flynn et al., 2020; Price et al., 2005). Then, candidate intact LTR-RTs were identified by LTR_retriever with default parameters (Combined with both LTR harvest and LTR_finder) (Li & Richard, 2009; Xu & Wang, 2007). The high-quality intact LTR-RTs were then produced by a custom program. We downloaded 7,497 *Gag-Pol* protein sequences of green plants from the National Center for Biotechnology Information (NCBI) as seed sequences and compared them with candidate intact LTR-RTs by Tblastn. If candidate LTR-RT contains the Gag-Pol sequence, it is confirmed as an intact LTR (Lyu et al., 2018). The LTR-RTs insertion time was estimated by LTR_retriever, and Matlab was used for graphic display. For solo- and truncated LTR-RTs, the non-redundant LTR library generated by LTR_retriever was compared with the whole genome sequences by Blastn. Three kb sequences of both upstream and downstream of every LTR paralog were extracted to compare with the 7,497 *Gag-Pol* protein sequences by Blastx. The LTR paralogs that lacked both upstream and downstream *Gag-Pol* sequences were considered as solo-LTRs (threshold setting with identity>30%, e-value <1e-8, and coverage >50%), those LTRs with *Gag-Pol* at only one side of flanking sequences were retained as truncated LTR-RTs.

### Gene family clustering analysis

OrthoFinder v2.5.2 was used for the gene family identification in *P. patens* and lycophytes, including *L. clavatum, I. taiwanensis, S. tamariscina*, and *S. moellendorffii* with parameters: -M msa -T raxml -S diamond (Buchfink et al., 2015; Emms & Kelly, 2019; Kozlov et al., 2019). The newly gained and lost gene families were inferred by dolloparsimony software (part of Tree2gd) with Dollo parsimony method.

### Analysis of the Huperzine A biosynthesis enzymes

We used the five HupA biosynthesis-related enzyme genes (*LDC, CAO, PKS, BBE*, and *SLS*) of *Huperzia serrata* (Yang et al., 2017) as references and compared them with the genomes of *L. clavatum* and other six species (*P. patens, I. taiwanensis, S. moellendorffii, S. tamariscina, S. lepidophylla*, and *A. thaliana*) by Blastp (with parameter: evalue <1e-5, identity >40 and score >100). Then we calculated the gene copy number of each HupA biosynthesis enzyme in different species by custom Perl script.

## ACKNOWLEDGEMENTS

This work was supported by the National Natural Science Foundation of China (Grant No.32170233, Grant No.32270248), and the Youth Innovation Promotion Association CAS (Grant No.2021075). We thank Xiang Zi-Yu for providing technical assistance for data analysis.

## AUTHOR CONTRIBUTIONS

Xiang QP and Zhang XC conceived the project, designed the experiments, and revised the manuscript. Yu JG and Tang JY wrote the manuscript. Yu JG and Lan MF analyzed the data. Yu JG developed the Pipeline for Post-assembly Decontamination. Yu JG and Tang JY finalized the figures and tables. Wei R and Xiang RC assisted in writing and editing. All authors have read and approved the manuscript.

## DATA AVAILABILITY STATEMENT

The whole genome sequence data reported in this paper have been deposited in the Genome Warehouse in National Genomics Data Center (Chen et al., 2021; CNCB-NGDC Members & Partners, 2022), Beijing Institute of Genomics, Chinese Academy of Sciences / China National Center for Bioinformation, under accession number GWHBJYW00000000 that is publicly accessible at https://ngdc.cncb.ac.cn/gwh. And the gene annotations are available at https://figshare.com/articles/dataset/Lycopodium_clavatum_genome_annotation/20493417.

## SUPPORTING INFORMATION

**Figure S1**. K-mer distribution of the *Lycopodium clavatum* genome. K-mer=17 Depth and K-mer number frequency distribution.

**Figure S2**. Non-redundant protein database (NR) homologous species distribution of the *Lycopodium clavatum* genome.

**Figure S3**. A cluster of Orthologous Groups of proteins (COG) functional classification of the *Lycopodium clavatum* genome.

**Figure S4**. Clusters of orthologous groups for eukaryotic complete genomes (KOG) functional classification of the *Lycopodium clavatum* genome.

**Figure S5**. Regression analyses of genome size on long terminal repeat (LTR) retrotransposons. **a**, Regression analysis of the number of intact LTR-RTs in four lycophyte genomes against assembled genome sizes. **b**, Regression analysis of the number of truncated LTR-RTs in four lycophyte genomes against assembled genome sizes. **c**, Regression analysis of the number of solo-LTR-RTs in four lycophyte genomes against assembled genome sizes. **d**, Regression analysis of the ratios of solo-LTR to intact LTR-RT (*S:I*) in four lycophyte genomes against assembled genome sizes.

**Figure S6**. Homologous dotplot within the *Lycopodium clavatum* genome.

**Figure S7**. Non-redundant protein database (NR) homologous species distribution of the *Lycopodium clavatum* genome.

**Table S1**. Summary of *Lycopodium clavatum* genome Illumina sequencing data. **Table S2**. Statistics of sequencing data obtained by Pacbio platform for *Lycopodium clavatum* genome.

**Table S3**. Genome survey analysis of *Lycopodium clavatum* obtained by Genome Characteristics Estimation (GCE).

**Table S4**. Illumina paired-end reads mapped to *Lycopodium clavatum* genome

**Table S5**. Analysis result of CEGMA v2.5.

**Table S6**. Analysis result of Benchmarking Universal Single-Copy.

**Table S7**. LTR Assembly Index score of *Lycopodium clavatum* genome.

**Table S8**. The statistics of repeat sequences in *Lycopodium clavatum* genome.

**Table S9**. The statistics of repeat sequences in *Lycopodium clavatum* genome.

**Table S10**. The statistics of NR Homologous Species Distribution.

**Table S11**. The go annotation in *Lycopodium clavatum* genome.

**Table S12**. The statistics of COG enrichment in *Lycopodium clavatum* genome.

**Table S13**. The statistics of KOG enrichment in *Lycopodium clavatum* genome.

**Table S14**. The statistic of RNA and pseudogene.

**Table S15**. The statistic of the GC content of all assembled contigs.

**Table S16**. The statistics of Blastn result between abnormal GC contig and NT database.

**Table S17**. GC content of different regions on abnormal GC.

**Table S18**. Statistics of non-plant sequences.

**Table S19**. The statistics of COG enrichment in non-plant sequence.

**Table S20**. The statistics of non-plant gene annotation by KEGG.

**Table S21**. The overall statistics of Orthofinder.

**Table S22**. The statistics of homologous family information of each species.

**Table S23**. The statistics of COG enrichment in *Lycopodium clavatum* unique genes.

**Table S24**. The statistics of KOG enrichment in *Lycopodium clavatum* unique genes.

**Table S25**. The statistics of LTR-RTs in lycophytes.

**Table S26**. The WGD gene pairs in *Lycopodium clavatum* by DupGen_finder.

**Table S27**. The dispersed gene pairs in *Lycopodium clavatum* by DupGen_finder.

**Table S28**. The statistics of low copy homologous gene trees by tree2gd.

**Table S29**. Top 2 of Blastp pairs and its Ks value.

**Table S30**. Kernel function analysis of Ks distribution.

**Table S31**. The copy numbers of five enzyme genes within HupA biosynthesis pathway in six species.

**Table S32**. The result of comparison between 500,000 random Illumina clean reads and NT database in *Lycopodium clavatum*.

**Table S33**. Reported HupA-producing fungi in non-plant sequences.

