## Supplementary figures and images for "The first homosporous lycophyte genome revealed the association between the dynamic accumulation of LTR-RTs and genome size variation"

### Figures S1-S7

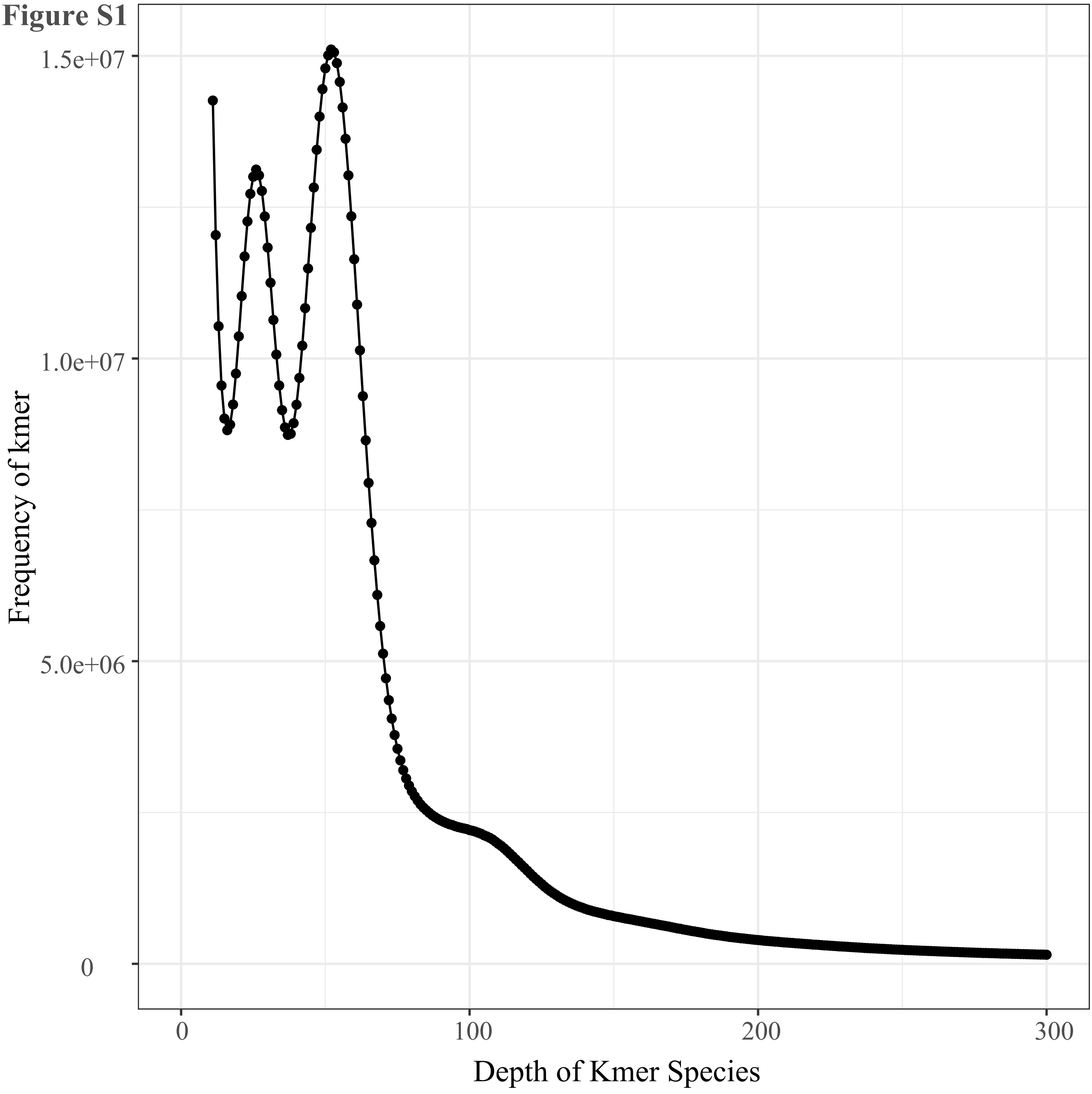

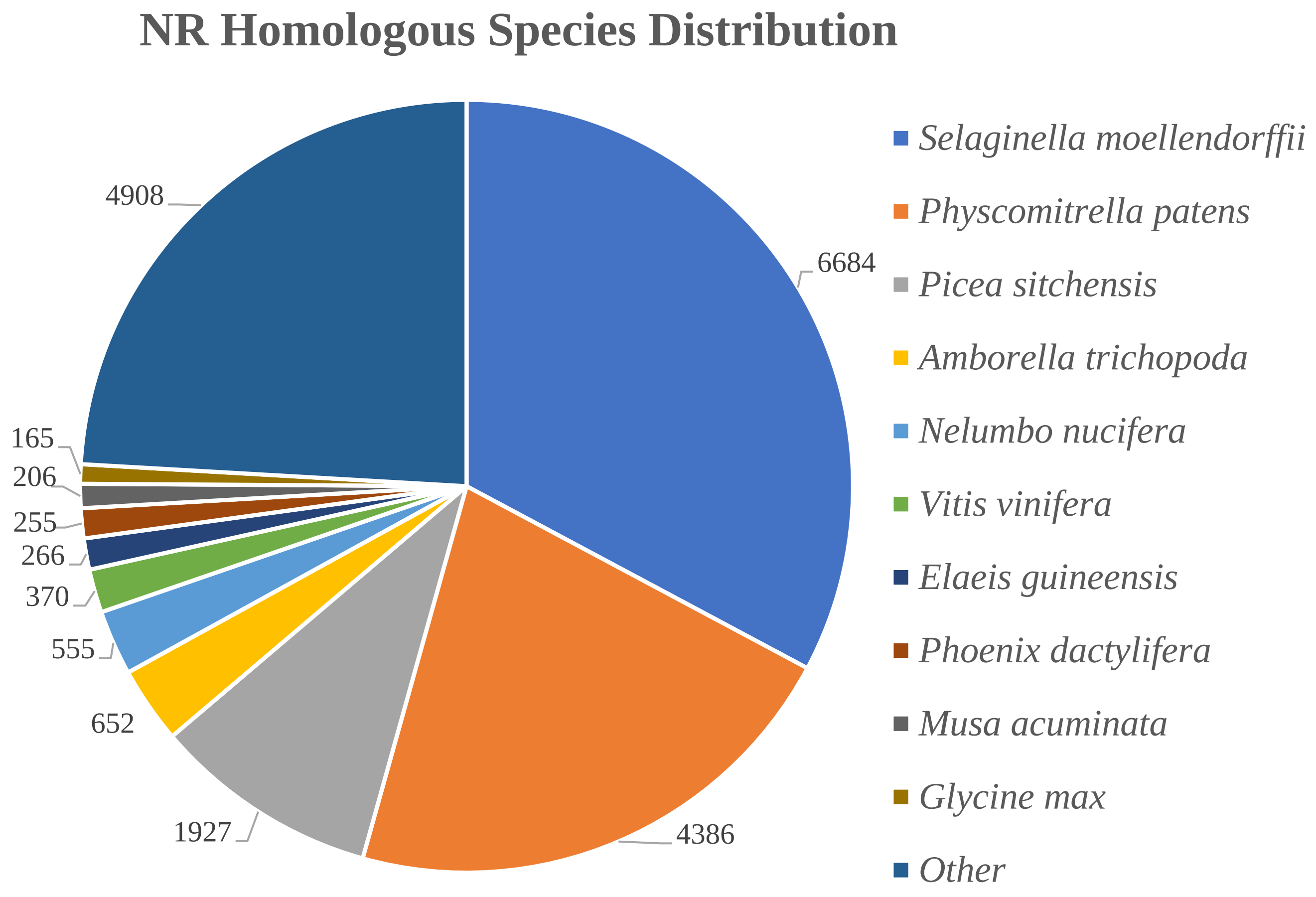

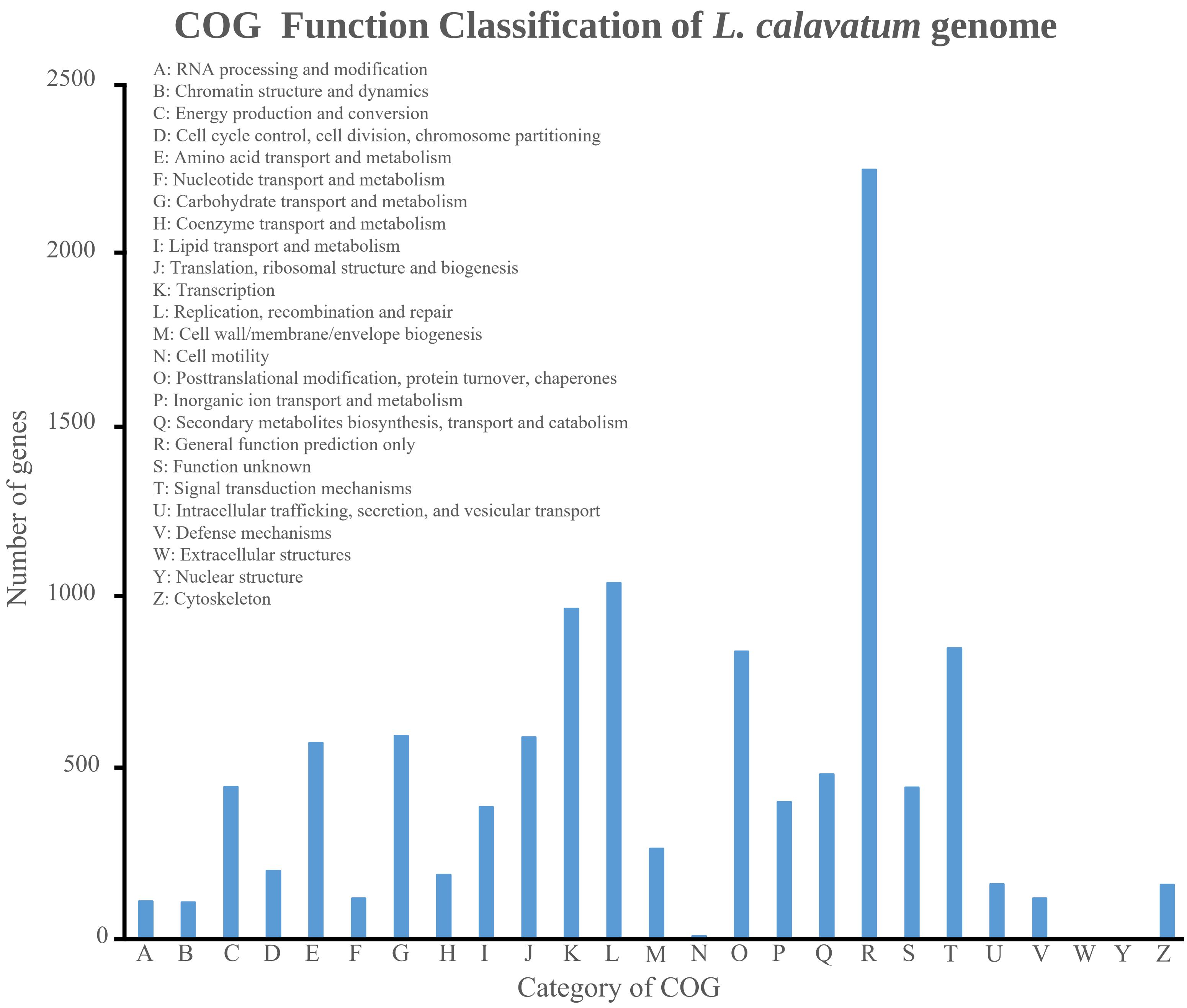

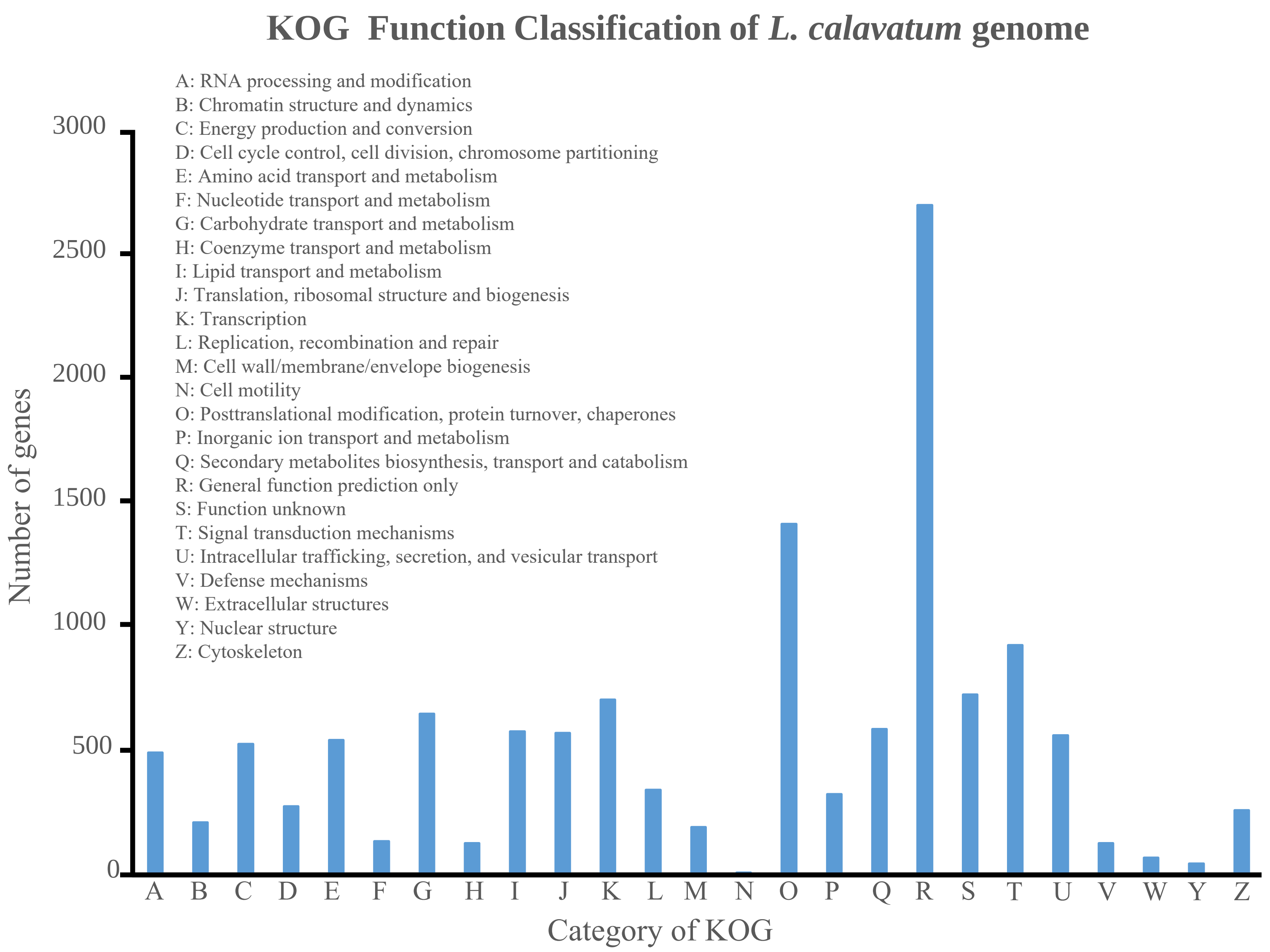

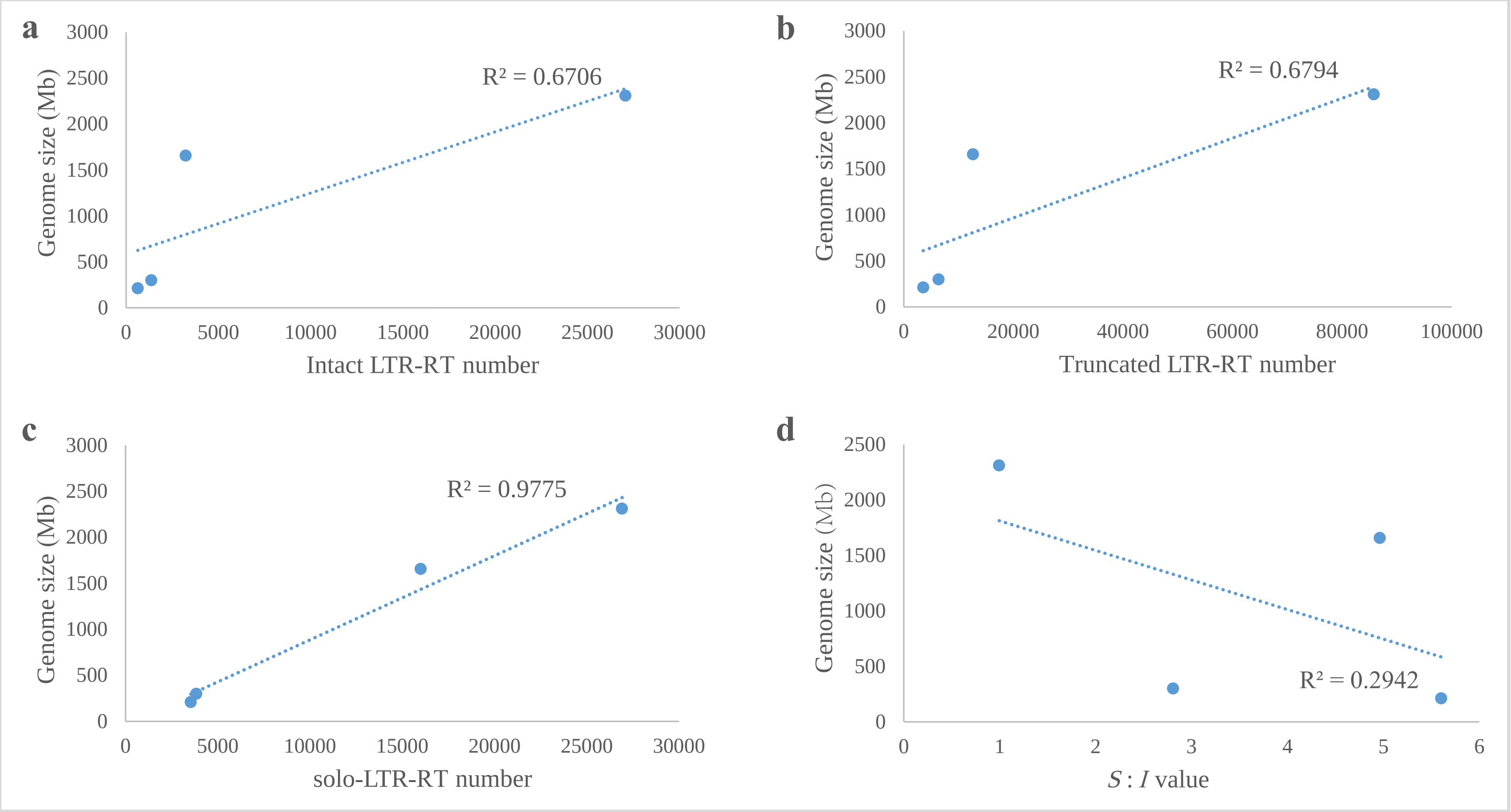

Figure S6

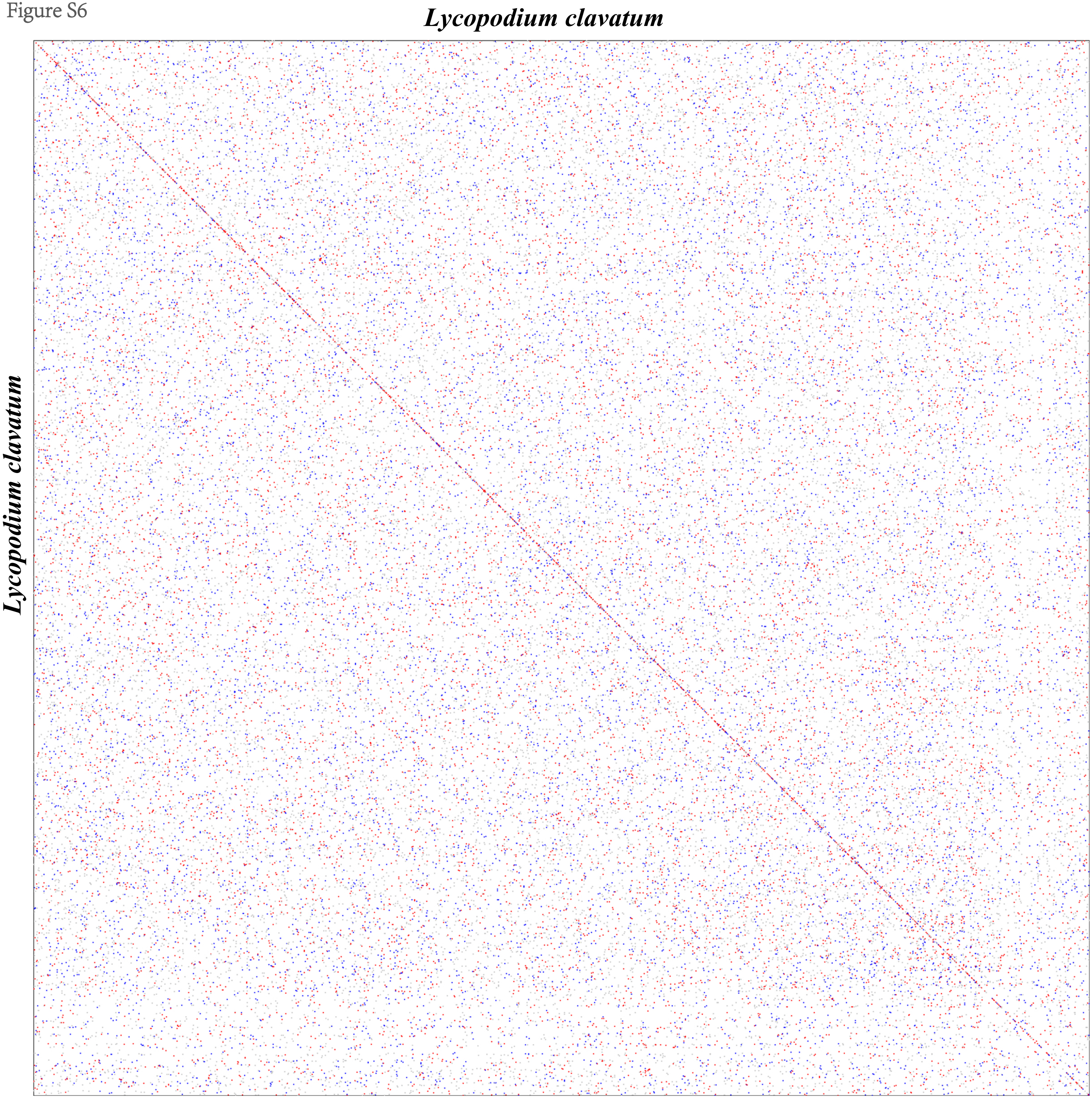

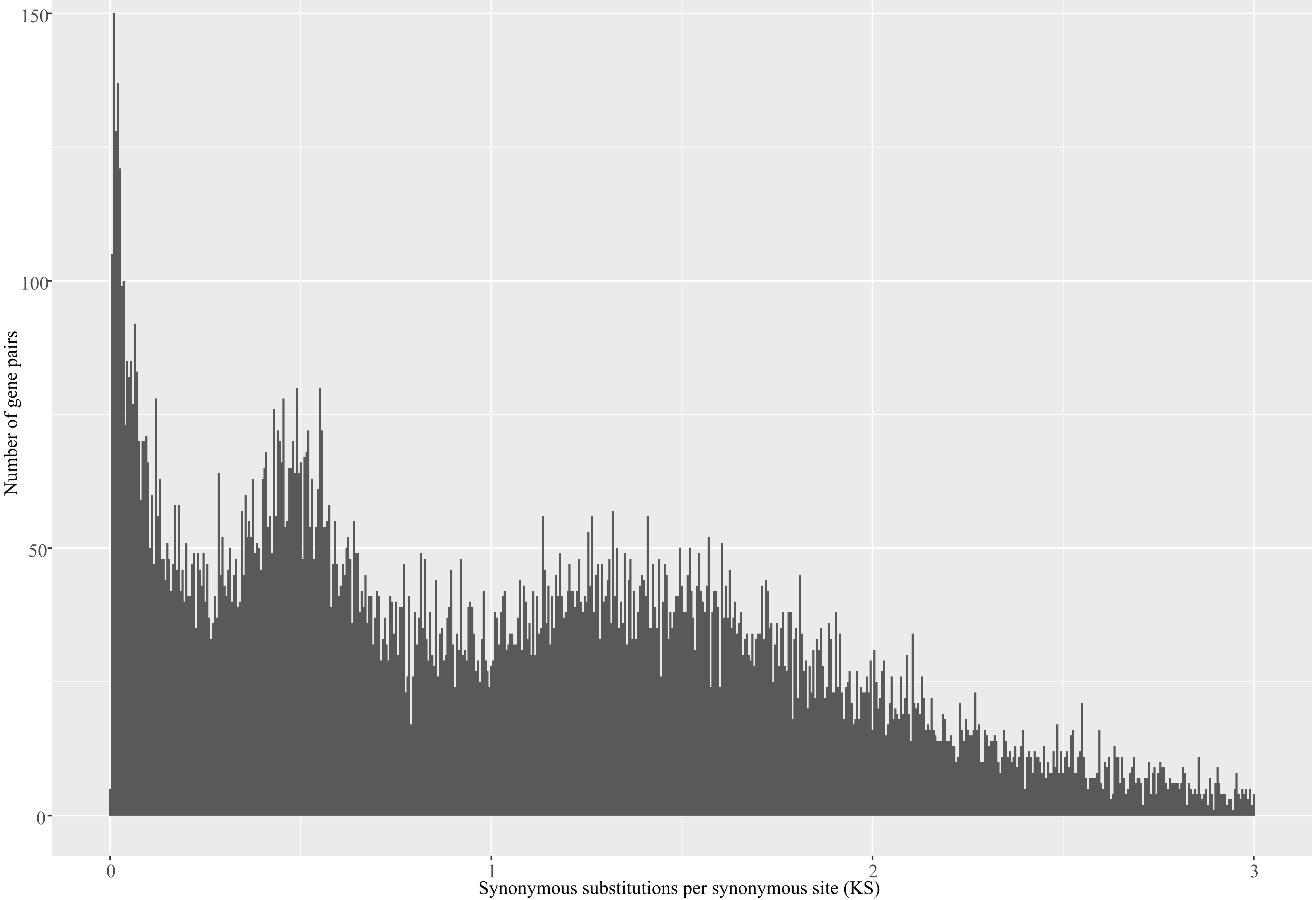
